# Dynamin-like proteins are essential for vesicle biogenesis in *Mycobacterium tuberculosis*

**DOI:** 10.1101/2020.01.14.906362

**Authors:** Shamba Gupta, Ainhoa Palacios, Atul Khataokar, Brian Weinrick, Jose L. Lavín, Leticia Sampedro, David Gil, Juan Anguita, M. Carmen Menendez, M. Jesus García, Navneet Dogra, Matthew B. Neiditch, Rafael Prados-Rosales, G. Marcela Rodríguez

## Abstract

*Mycobacterium tuberculosis (Mtb)* secretes pathogenicity factors and immunologically active molecules via membrane vesicles. However, nothing is known about the mechanisms involved in mycobacterial vesicle biogenesis. This study investigates molecular determinants of membrane vesicle production in *Mtb* by analyzing *Mtb* cells under conditions of high vesicle production: iron limitation and VirR restriction. Ultrastructural analysis showed extensive cell envelope restructuring in association with vesicle release that correlated with downregulation of cell surface lipid biosynthesis and peptidoglycan alterations. Comparative transcriptomics showed common upregulation of the *iniBAC* operon in association with high vesicle production in *Mtb* cells. Vesicle production analysis demonstrated that the dynamin-like proteins (DLPs) encoded by this operon, IniA and IniC, are necessary for release of EV by *Mtb* in culture and in infected macrophages. Isoniazid, a first-line antibiotic, used in tuberculosis treatment, was found to stimulate vesicle release in a DLP-dependent manner. Our results provide a new understanding of the function of mycobacterial DLPs and mechanistic insights into vesicle biogenesis. The findings will enable further understanding of the relevance of *Mtb*-derived extracellular vesicles in the pathogenesis of tuberculosis and may open new avenues for therapeutic research.

**IMPORTANCE:** Iron is an essential nutrient that promotes survival and growth of *M. tuberculosis*, the bacterium that causes human tuberculosis (TB). Limited availability of iron, often encountered in the host environment, stimulates *M. tuberculosis* to secrete membrane-bound extracellular vesicles containing molecules that may help it evade the immune system. Characterizing the bacterial factors and mechanisms involved in the production of mycobacterial vesicles is important for envisioning ways to interfere with this process. Here, we report the discovery of proteins required by *M. tuberculosis* for vesicle biogenesis in culture and during host cell infection. We also demonstrate a connection between antibiotic response and extracellular vesicle production. The work provides insights into the mechanisms underlying vesicle biogenesis in *M. tuberculosis* and permits better understanding of the significance of vesicle production to *M. tuberculosis*-host interactions and antibiotic stress response.

## INTRODUCTION

Extracellular vesicles (EV) are membrane-bound structures actively secreted by most, if not all, cells and play important roles in intercellular communication. Bacteria release EV containing proteins, genetic material, and lipids to communicate with both prokaryotic and eukaryotic cells in their environment (1, 2).

*M. tuberculosis* (*Mtb*), the causative agent of human tuberculosis (TB), releases EV during axenic growth and into the macrophages it infects, both in culture and in the lungs of infected mice (3). *Mtb*-produced EV (*Mtb*-EV) contain many immunologically active lipoproteins and glycolipids (3, 4) and are enriched in polar lipids consistent with a cytoplasmic membrane origin (3). *Mtb*-EV mediate transfer of bacterial components into immune cells (5, 6) and aid in acquisition of nutritional iron (Fe) by mediating export and uptake of the hydrophobic siderophore, mycobactin (7). Furthermore, *Mtb*-EV contain protective antigens; immunization with isolated EV elicits a protective immune response against TB in mice that is comparable to BCG vaccination (8).

Although these observations suggest *Mtb*-EV play a role in TB pathogenesis and immunity, current knowledge about the mechanistic and molecular aspects of mycobacterial EV biogenesis is very limited. Vesicle biogenesis is known to be an active and genetically regulated process that can be triggered by iron deprivation (3, 7). EV production is also promoted by loss of the *rv0431* product, VirR (Vesiculogenesis and immune <u>r</u>esponse <u>R</u>egulator), a member of the LytR-CpsA-Psr (LCP) protein family (9). This family includes enzymes that transfer glycopolymers from membrane-linked precursors to peptidoglycan or cell envelope proteins and are central to cell envelope integrity (10), but the function of VirR is unknown.

In this study, we investigate molecular determinants of vesicle release by examining common cellular and gene expression features of hypervesiculating *Mtb*. We report that production of EV is associated with extensive restructuring of the cell envelope compatible with regulation of outer membrane biogenesis in Fe-limited mycobacteria and altered peptidoglycan dynamics in the *Mtb virR* mutant (*virR*). We show that the induction of the *iniBAC* operon is common to the transcriptional response of *virR* and Fe-deprived *Mtb*. This operon is well recognized as a marker of the response to cell envelope biogenesis targeting antibiotics (11). Here, we demonstrate that the dynamin like proteins (DLPs) IniA and IniC are required for *Mtb*-EV biogenesis in culture and during macrophage infection. Further, we report that sub-inhibitory concentrations of the cell envelope active antibiotic isoniazid, which induces *iniBAC*, promote EV release. Our results have uncovered molecular determinants of EV production in mycobacteria, opening the way to a better understanding of the mechanisms of vesicle biogenesis and the relevance of vesicle mediated immunomodulation in TB pathogenesis.

## RESULTS

### High EV production is associated with an altered cell envelope in *Mtb*

While Gram-negative bacteria can release outer membrane vesicles directly into the extracellular environment, mycobacteria have a thick cell envelope surrounding the cytoplasmic membrane that acts as a permeability barrier. This cell envelope is composed of peptidoglycan (PG) covalently linked to arabinogalactan (AG), forming the cell wall, which in turn is decorated with exceptionally long-chain mycolic acids (MA). These MA, together with intercalating glycolipids, form the unique mycobacterial outer membrane (OM) (12, 13).

Analysis of vesicle associated lipids showed predominantly polar lipids, consistent with the cytoplasmic membrane being the likely origin of the vesicles (3). This is true also in EV produced by Fe-limited *Mtb* whose lipid content consist mainly of polar lipids and the lipidic siderophore, mycobactin (7). Given that these vesicles must traffic from their point of origin in the plasma membrane through the cell envelope, we hypothesize that local remodeling of the cell envelope is necessary for EV release, and that this process may be exacerbated in a *virR* mutant and Fe-deprived *Mtb*. To test this, we examined the cell envelope structure of a transposon insertion mutant of *rv0431* (*virR*) (9) grown in high Fe medium, as well as wild type (WT) *Mtb* grown in high and low Fe conditions (Fig. 1). Using cryo-electron microscopy (Cryo-EM) on whole cells, which allows the study of mycobacterial cell surface-associated compartments in a close-to-native state (13, 14), we could easily discern the three layers comprising the mycobacterial cell envelope: i) the outer membrane (OM); ii) an intermediate layer composed of two layers (L1 and L2, previously assigned to the mycolate-PG-AG (mAGP) network) (13, 14)); and iii) the cytoplasmic membrane (CM) (Fig. 1). We observed a significant difference in cell envelope thickness (OM-CM) in the *virR* mutant strain compared to the WT. This difference was largely accounted by expansion of the CM-L1 layer suggesting cell wall alterations. Complementation with a functional copy of *virR* restored normal cell envelope structure (Fig. 1A-C, E). In Fe-deprived *Mtb*, we observed a significant reduction in the thickness of the OM in comparison to Fe-sufficient cells (Fig 1D-E). These changes in the cell envelope of high EV producing bacteria suggest that cell envelope alterations in response to Fe-deficiency or lack of VirR and increased membrane vesicle biogenesis might be connected in *Mtb*.

**Figure 1.**
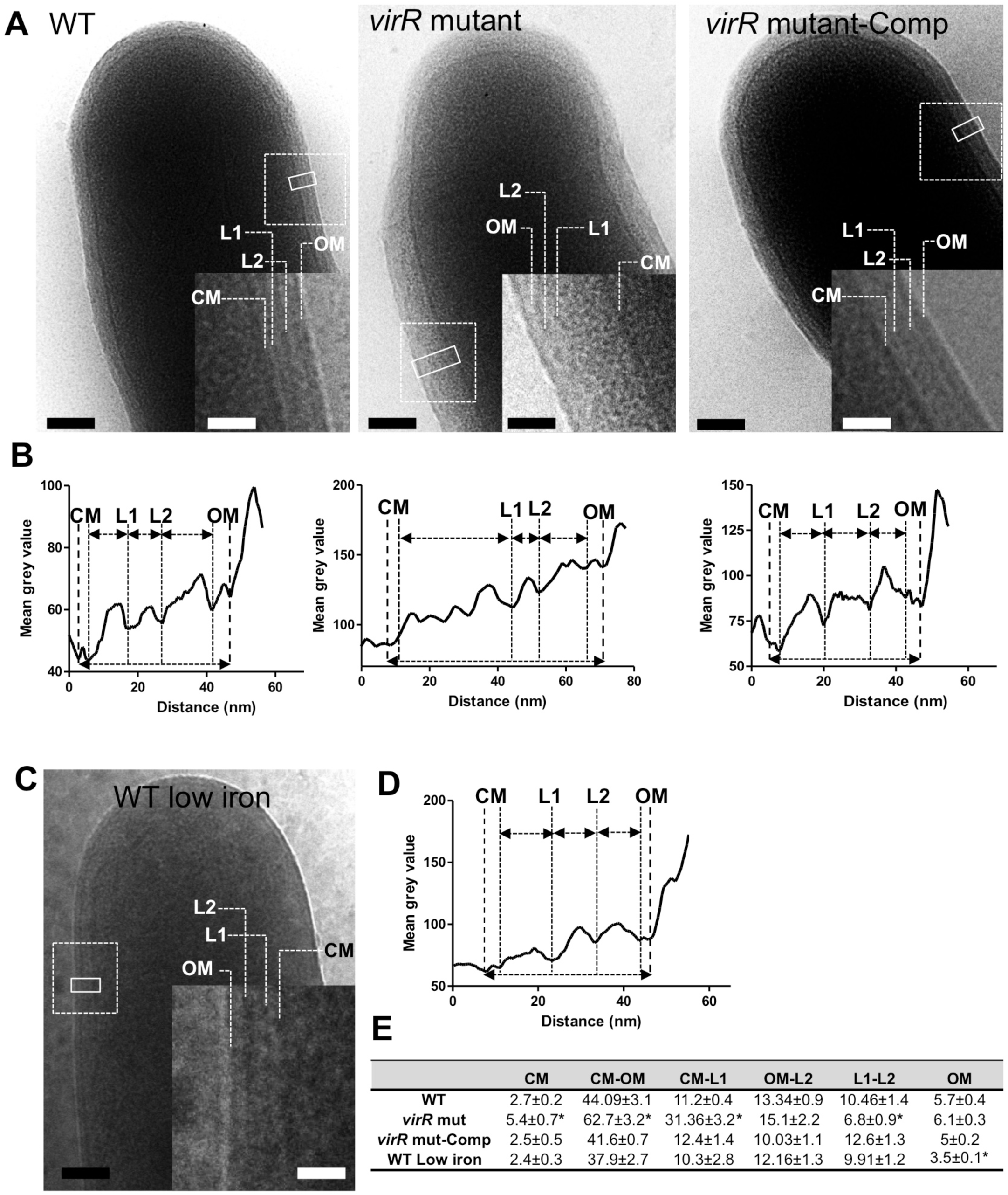
Ultrastructural changes in the cell envelope of *virR* and Fe deprived *Mtb* associated with increased vesiculation. (**A**) Cryo-electron micrographs of indicated *Mtb* strains grown in high Fe MM. Closed line rectangles were used to calculate grey value profiles of cell envelopes using ImageJ. The dashed line insets within the main micrograph were magnified to show a detailed view of the cell surface. Scale bars are 100 nm in main micrographs and 50 nm in the insets. (**B**) Density profiles based on grey values of the cross sections marked by solid line rectangles in A. (**C**) Cryo-EM micrograph of *Mtb* grown in low Fe MM. (**D**) Density profile based on grey values of the cross section marked by the closed rectangle in C. (**E**) Mean values and standard errors of distances in nm between main cell envelope layers measured in A and C. \**P* < 0.05. CM, cytoplasmic membrane; OM, outer membrane; L1, layer 1; L2, layer 2. Images and measurements shown are representative of 50 independent cells.

To better understand this relationship, we reanalyzed our published transcriptomic data comparing expression of cell envelope genes in Fe-deprived and Fe-sufficient *Mtb* (15). Iron-deprived bacteria showed repression of genes involved in MA and dimycocerosate ester (DIM) biosynthesis, as well as the Ag85 complex which transfers a mycolate chain to AG polysacharides to form the mAGP complex and build the OM (Fig. S1). These results are consistent with biochemical analysis of Fe-deprived mycobacterial cells that showed reduction of surface mycolates and enhanced cell envelope permeability (16, 17). Additionally, we found the ultrastructural changes observed in the *virR* mutant correlated with increased susceptibility to the muramidase lysozyme and increased release of muropeptides relative to WT (Fig. S2), indicating modifications in PG remodeling in VirR restricted cells. Collectively, these observations suggest the possibility that structural changes associated with down modulation of cell envelope lipid biosynthesis during Fe-limitation, and alterations in VirR-regulated PG dynamics support increased release of EV in *Mtb*.

### Identification of candidate vesicle biogenesis genes

To identify mediators of vesicle production, we first carried out transcriptional profiling of *virR* and WT *Mtb*. Although previous efforts to define the transcriptional program of *virR* showed no significant differences in gene expression compared to WT (9), those experiments were carried out with bacteria grown in rich 7H9 medium. Here, we instead used a defined high-Fe minimal medium (MM) (3) and were able to identify 65 downregulated and 15 upregulated genes in *virR* compared to WT (Fig. 2). Genes associated with hypoxic response, metal homeostasis (*csoR, cmtR, kmtR, ctpS*), carbon and nitrogen starvation (*lat, cysD, usfY, sigE)*, heat shock, and lipid biosynthesis were found expressed at lower levels in *virR* than in WT (Fig. 2B). Conversely, gene transcripts associated with response to isoniazid (*iniBAC*), PDIM biosynthesis, and Fe-S clusters [Fe-S] biosynthesis were more abundant in *virR* (Fig. 2*B*). Notably, the entire dormancy survival regulon (DosR) was downregulated in the mutant (Fig. 2C) suggesting the mutant is less sensitive to signals that activate the DosR response, which include oxygen and nutrient deficiency (18).

**Figure 2.**
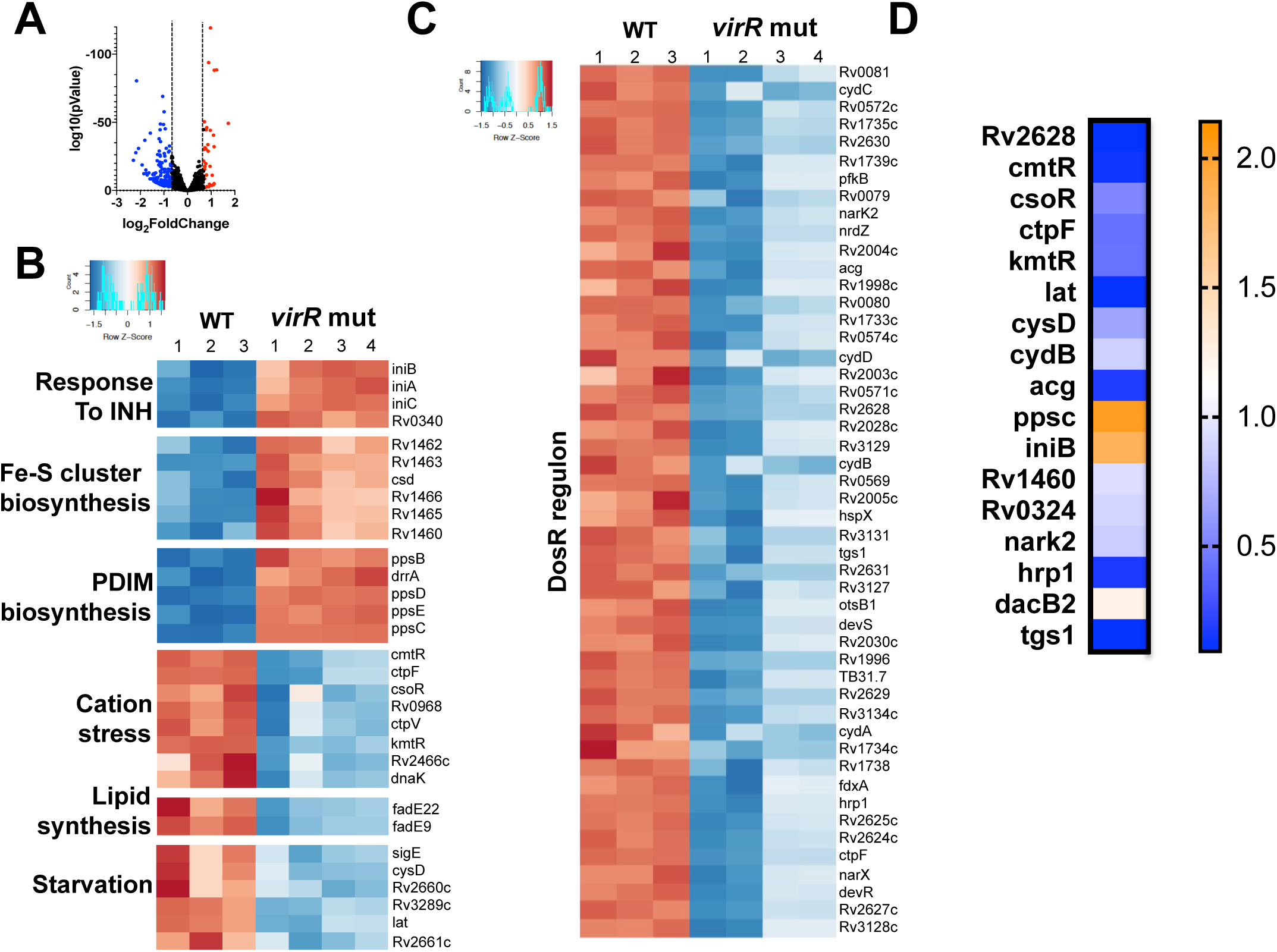
Transcriptional signature of the *virR* mutant strain. (**A**) Volcano plots showing the genes upregulated (red) and downregulated (blue) in the *virR* strain relative to the WT strain. Expression threshold was set at log_2_ fold change of +/- 0.68. (**B**) Heatmap representing gene expression in *virR* and WT *Mtb*. Genes are grouped according to different physiological responses or biosynthetic pathways. (**C**) Heatmap showing levels of expression for genes belonging to the DosR/T regulon in *virR* and WT *Mtb*. Numbering (1, 2, 3 and 4) denotes the independent biological replicates. The intensity of the colors is determined by the calculated *Z* value for each condition. (**D**) Validation of selected genes in the RNAseq data from *virR* by quantitative RT-PCR.

Next, we compared the transcriptomics data from the *virR* mutant to our published transcriptional profiling data from iron-deprived *Mtb* (15) with the hypothesis that molecular determinants of vesicle biogenesis are induced in both hyperversiculation conditions. We found that the *iniBAC* operon and [Fe-S] assembly genes were induced in both *virR* and iron-deprived *Mtb*, relative to WT *Mtb* grown in high-iron (Fig. S3). Notably, the growth of *virR* mutant cells in low-iron medium is slower than WT (Fig. S4), which suggests that increased sensitivity to iron limitation underlies the enhanced expression of [Fe-S] assembly genes. We also found genes downregulated in *virR* and upregulated in iron-deprived *Mtb*, such as *rv0079*-*rv0081* (encoding a hypoxia response transcriptional regulator), genes encoding for universal stress proteins (*rv1996, hspX, rv2624c, hsp1*), and genes involved in the propionate assimilation pathway (*prpCD*) and triacylglycerol synthesis (*tgs1*). These findings indicate that the metabolic stresses generated by *virR* inactivation and Fe limitation, are not identical, supporting the notion that genes upregulated in both conditions reflect convergence upon a shared function, such as vesicle biogenesis.

Previous reports support a link between cell envelope stress and induction of the *iniBAC* operon: the *iniBAC* operon was originally identified as specifically induced by the antibiotics isoniazid (INH) and ethambutol, both of which inhibit cell envelope synthesis (11, 19) and recently, induction of *iniBAC* has been linked to the biosynthesis of methyl-branched lipids, (20). While IniB is homologous to an *Arabidopsis thaliana* cell wall protein of unknown function, *iniA* and *iniC* encode GTP-binding domain containing proteins related to dynamin-like proteins (DLPs) (Fig. 3A) (21).

**Figure 3.**
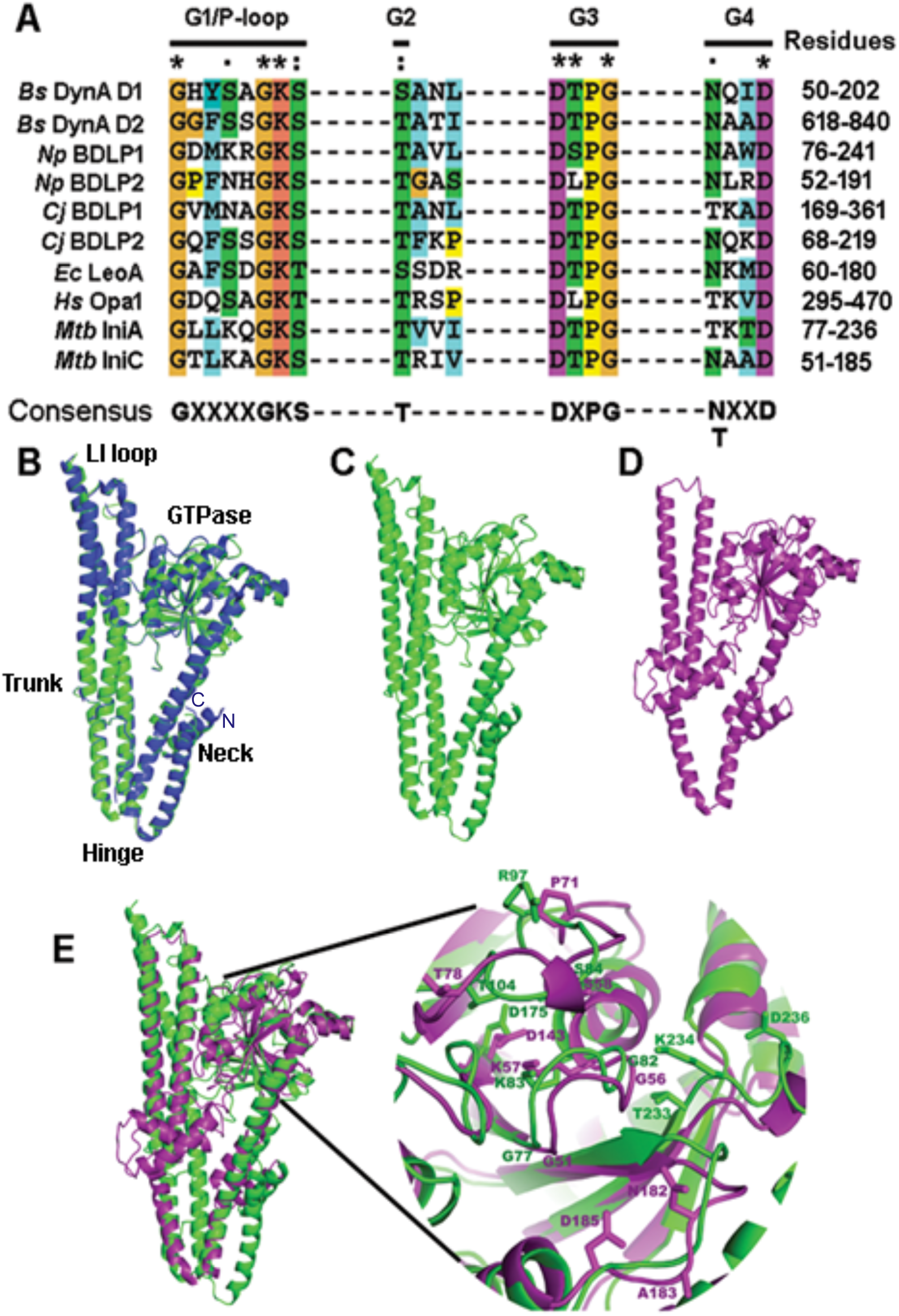
Sequence and structural alignment of mycobacterial DLPs. (**A**) Sequence alignment of *Mtb* IniA and IniC G1-G4 GTP binding motifs with other members of the dynamin family. *Bacilllus subtilis* DynA (P54159), *Nostoc punctiforme* BDLP1 (B21ZD3) and BDLP2 (B21ZD2), *Campylobacter jejuni BDLP1 (*CJ0411) and BDL2 (CJ0412), *Escherichia coli* LeoA (E3PN25) and human mitochondrial OPA1 (0060313). (**B**) Computational model of *Mtb*-IniA (green) superimposed on structure of *Msm*-IniA (blue, PDB id: 6J73). (**C**) Computational model of *Mtb*-IniA generated from the crystal structure of *Msm*-IniA (PDB id: 6J73). (**D**) Computational model of *Mtb*-IniC generated from the crystal structure of *Msm*-IniA (PDB id: 6J73). (**E**) Model of *Mtb*-IniC (purple) superimposed on model of *Mtb*-IniA (green) zooming in the GTPase domains of IniA and IniC showing the residues proposed to be involved in GTPase activity in IniA and IniC.

Classic dynamin and DLPs form a large family of GTPases that mediate membrane fission and fusion in both eukaryotic and prokaryotic cells. Although the IniA protein in *Mtb* has not been functionally characterized, the crystal-structure of *M. smegmatis* (*Msm*) IniA showed conserved structural features with other DLPs (22) including a GTPase domain containing the four consensus elements (G1-G4) and two elongated alpha-helical bundle domains which are structurally contiguous and are also known as neck and trunk domains (Fig 3B). *Msm*-IniA exhibits GTPase-dependent membrane fission activity *in vitro*, and it localizes to the plasma membrane causing cell surface wrinkling, when overexpressed in *Msm* (22). Computer modeling of *Mtb*-IniA based on the crystal structure of *Msm*-IniA suggests conserved folding between the two homologs (Fig 3B-C), although the annotated *Mtb* protein has an N-terminal extension of 34 amino acids. At the trunk tip, *Msm-*IniA has a lipid-binding domain composed of hydrophobic and positively charged residues that are necessary for lipid binding and membrane association (22). These residues are similar in *Mtb*-IniA. IniA and IniC share 21% sequence identity. IniC is a smaller protein. However, computational modeling suggests that like IniA, IniC has a core DLP-like fold with conserved GTPase, neck and trunk domains (Fig 3C-E). Together with our data showing that *iniBAC* expression is induced in hypervesiculation conditions, the similarity of IniA and IniC to DLPs, whose canonical role is in membrane restructuring supports IniA and IniC function in mycobacterial EV biogenesis.

### *iniAC* are necessary for efficient EV release in *Mtb*

To investigate the role of *iniBAC* genes in *Mtb*-EV biogenesis, we assayed EV production in a *Mtb* mutant in which *iniA* was replaced by a hygromycin-resistance cassette (*iniA*∷*hyg*) (23). EV were isolated from cultured *Mtb* strains grown in low Fe, and their abundance assessed via multiple parameters including quantification of protein and lipid content, nanoparticle analysis and semi-quantitative dot blot using a mouse antiserum previously raised against EV. All of these measurements showed drastic reduction in EV production in the *iniA∷hyg Mtb* mutant compared to the WT (Fig. 4 and Fig. S5)

**Figure 4.**
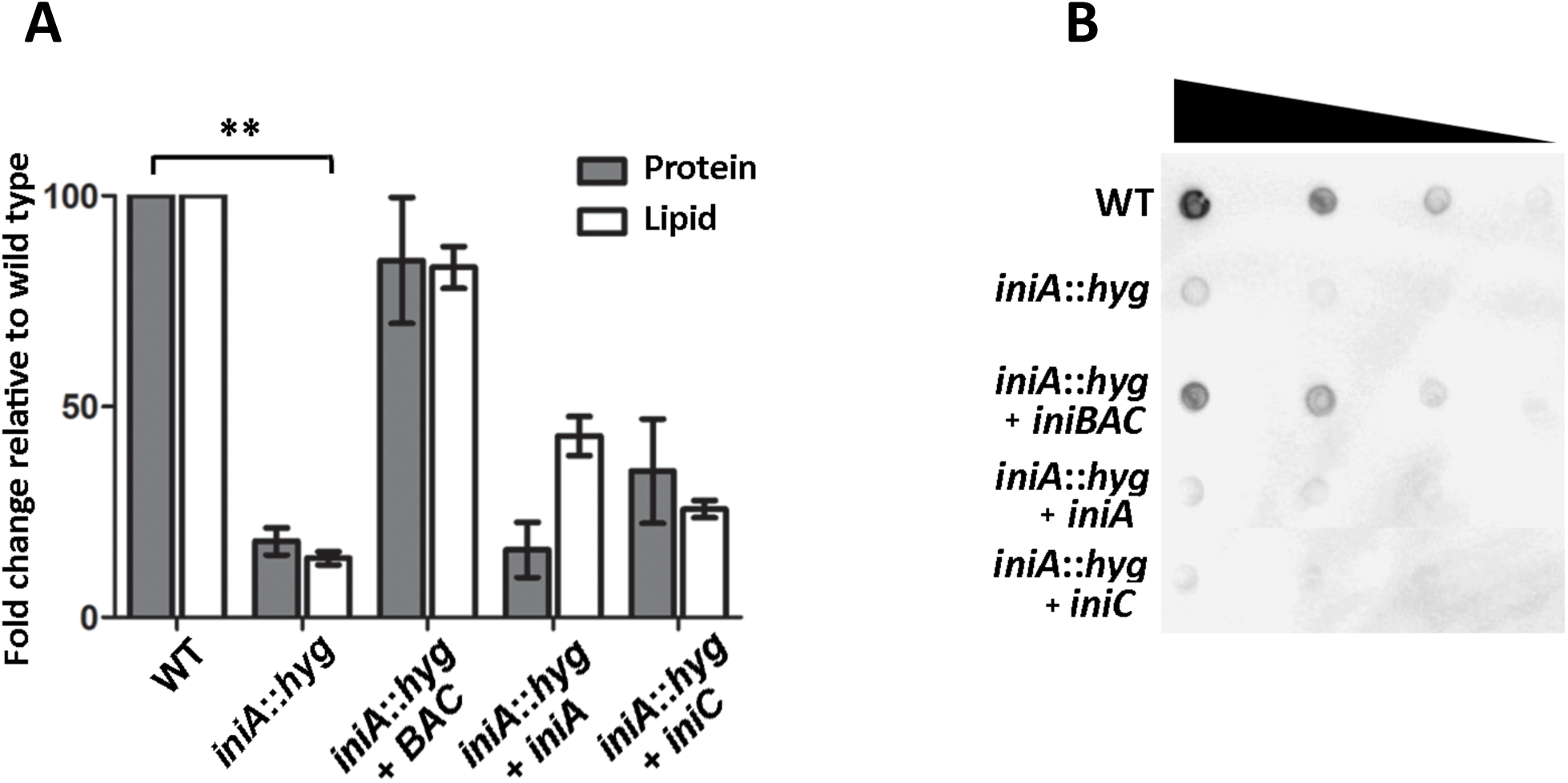
Release of *Mtb*-EV is dependent on DLPs in *Mtb*. (**A**). Quantification of EV production based on protein and lipid associated with *Mtb*-EV isolates purified by differential centrifugation and density gradient (as shown in Fig S5) and normalized to number of CFUs derived from indicated strains grown in low Fe MM. Data represents the mean ± SD of triplicate experiments \*\**P* < 0.005. (**B**) Dot blot of EV preparations isolated from indicated strains recognized by anti-EV polyclonal murine serum. The results shown are representative from 5 independent experiments.

The overall concentration of secreted protein in WT and *iniA∷hyg* culture supernatants was comparable, indicating *iniA∷hyg* does not have a pleotropic defect on protein synthesis or secretion (Fig. S6).

The defect in EV production observed in the *iniA∷hyg* mutant was to a large extent complemented by a wild-type copy of *iniBAC*, expressed under its native promoter in an integrative plasmid (Fig. 4), but was not complemented by *iniA* or *iniC* individually, indicating the *iniA* mutation is polar on *iniC* and that both IniA and IniC are necessary for vesicle production. Collectively the results demonstrate DLPs are required for EV biogenesis in *Mtb* and might work together in this process.

### Isoniazid treatment promotes EV production

Isoniazid (INH) is a first-line anti-TB drug that targets cell envelope biogenesis specifically mycolic acid biosynthesis. Since induction of the *iniBAC* operon is a hallmark of the mycobacterial response to INH we determined whether INH could trigger EV production in *Mtb*. We treated iron-sufficient cultures of *Mtb* with sub-inhibitory concentrations of antibiotic for five days and then estimated abundance of *Mtb*-EV in culture filtrates. Enhanced EV release was observed by dot blot analysis at the lowest concentration tested, 0.005 µg.ml^-1^ (Fig 5B). Quantification of protein and lipid associated with vesicles, indicated over 2-fold increased vesicle release at the highest INH concentration tested 50 ng/ml which is still 10 fold lower than the minimum inhibitory concentration (MIC) of INH in MM (24). This increase in EV release correlated with upregulation of *iniA* detected by RT-PCR (Fig. S7) and with visible cell surface changes, such as the formation of blebs and filaments in the INH-treated cultures, a phenotype we previously observed in the *virR* mutant (Fig. 5*C*) (9).

**Figure 5.**
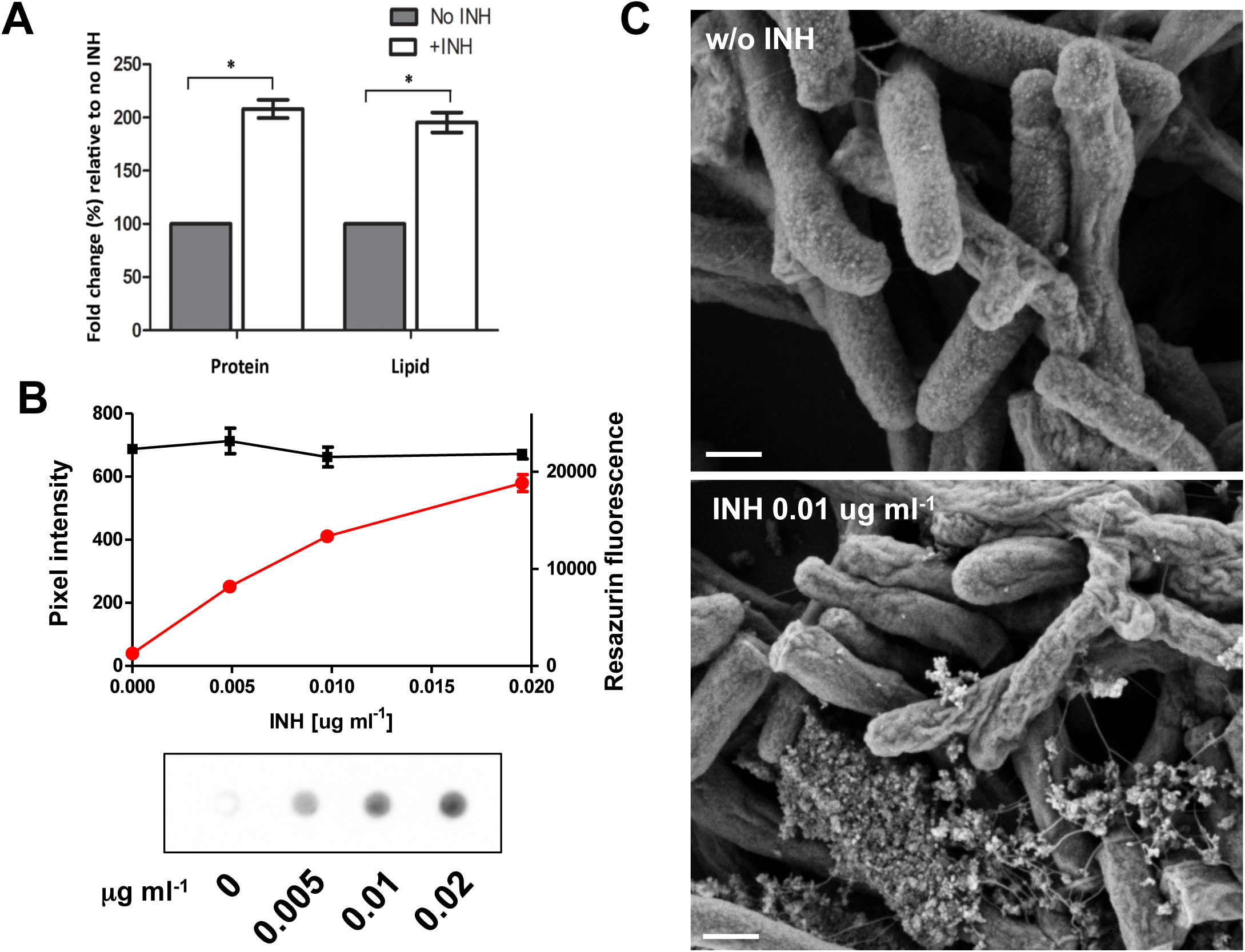
EV production in cells exposed to INH. (**A**) Quantification of protein and lipid associated with EV isolated from culture filtrates from WT grown in high Fe MM with 0 or 50 ng.mL^-1^ of INH normalized to CFUs. Data represents the mean ± SD of triplicate experiments. (**B**) WT *Mtb* cultured in high iron MM was subjected to sub-inhibitory concentrations of INH. For each concentration of antibiotic, cell viability was assessed by the resazurin cell viability assay (black line) and EV release was determined by dot-blot using an anti-EV polyclonal murine serum. The intensity of the dot-blot signal was determined by Image J and showed in the red line. (**C**) SEM micrographs of WT *Mtb* exposed to 0 or 10 ng.mL^-1^ INH. Scale bar corresponds to 200 nm. \**P* < 0.05

INH-induced EV release was abrogated in the *iniA∷hyg* mutant, independent of iron levels in the medium (Fig. 6), indicating that EV release in response to INH is dependent on DLPs. These findings suggest that sub-inhibitory concentrations of INH trigger *Mtb*-EV release via induction of *iniBAC*.

**Figure 6.**
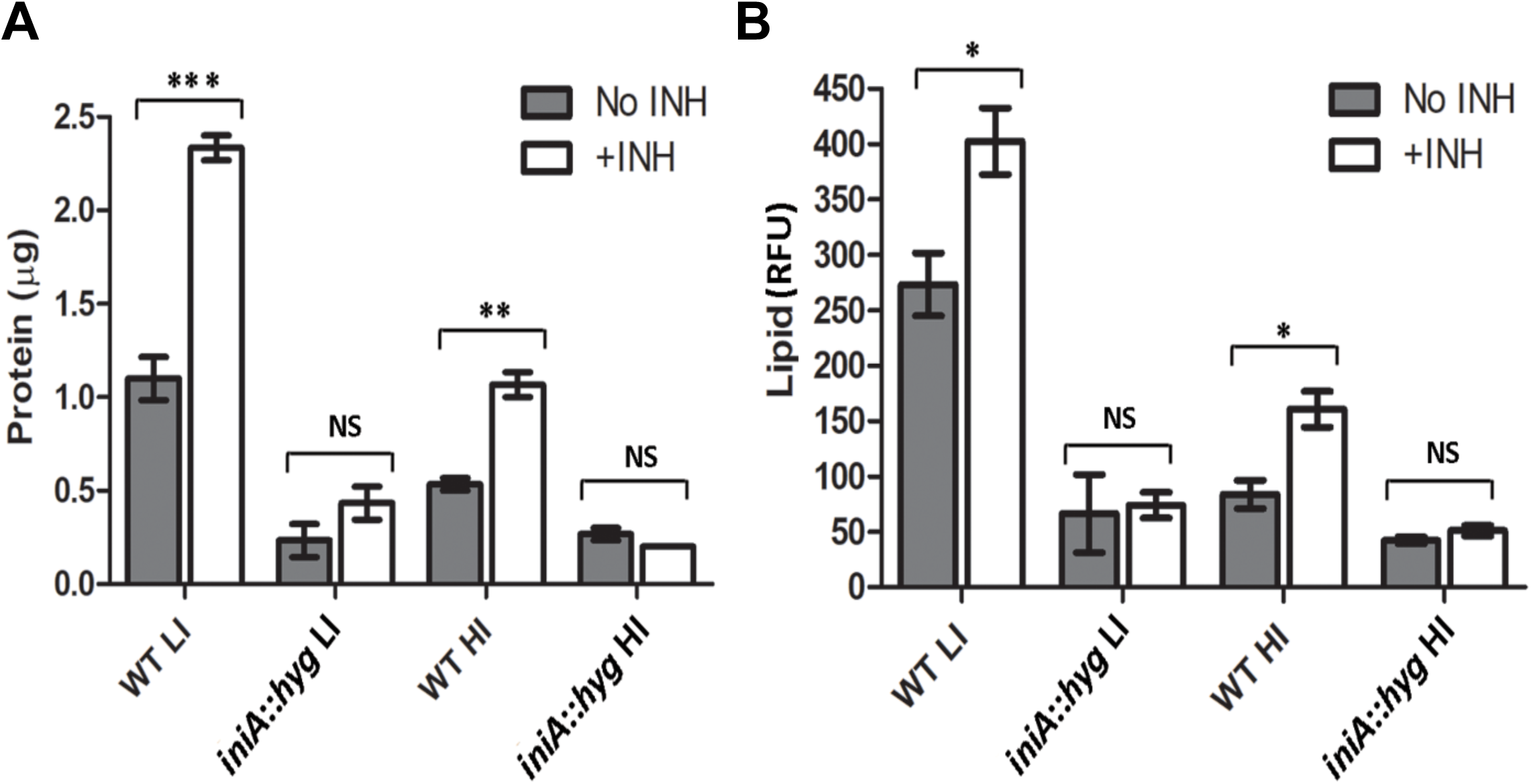
EV release in WT and *iniA∷hyg* exposed to INH. (**A**) Protein and (**B**) lipid associated with EV isolates from WT and *iniA∷hyg* cultured in low (LI) or high (HI) Fe MM with 0 or 50 ng.mL^-1^ INH. Data represents the mean ± SD of three independent experiments. \**P* < 0.05.

### Lack of DLPs abrogates vesicle-mediated iron acquisition in *Mtb*

When faced with iron limitation, *Mtb* produces mycobactin and carboxymycobactin, hydrophobic and amphiphilic siderophores, respectively. While carboxymycobactin is actively secreted via membrane transporters (25), mycobactin is localized in close proximity to the cytoplasmic membrane (26), and is exported via EV (7). Mycobactin containing EV can serve as the sole Fe source for Fe deficient *Mtb* (7). Accordingly, culture filtrates from the EV-deficient *iniA∷hyg* strain were unable to support the growth of a siderophore synthesis deficient strain of *Mtb* (ST142) (27) while culture filtrates derived from the equivalent number of WT *Mtb* cells were highly efficient (Fig 7A). This phenotype was complemented by a wild-type copy of *iniBAC* (Fig. 7A). Notably, the growth rate (Fig. 7B) and levels of mycobactin synthesized by *iniA∷hyg* and the WT *Mtb* in low Fe conditions were equal (Fig 7C), indicating that the reduced EV production in the absence of DLPs is not due to a defect in sensing and responding to Fe limitation. It also indicates that iron obtained through carboxymycobactin maintains normal growth of *iniA∷hyg* in culture medium. These results suggest that by abrogating EV production, the disruption of mycobacterial DLP function decreases availability of a public good *i*.*e*., extracellular mycobactin and may limit acquisition of Fe in the host, particularly in microenvironments where carboxymycobactin has restricted access to Fe or is sequestered by the host siderophore binding protein, lipocalin (28) which is abundant in TB granulomas (15).

**Figure 7.**
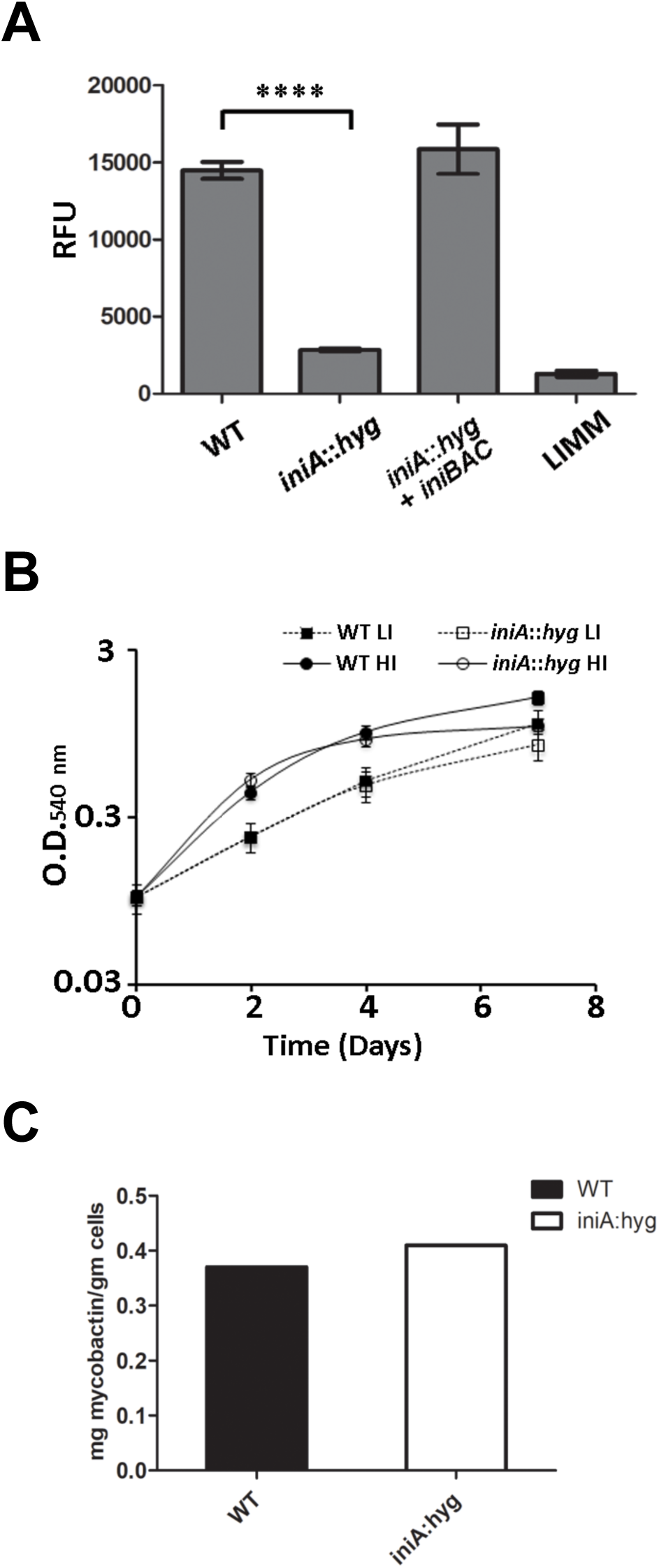
EV-mediated Fe acquisition is dependent on DLPs. (**A**). The iron uptake deficient *mbtB* mutant of *Mtb* (ST142) was grown in MM low iron (LIMM) or LIMM supplemented with EV preparations derived from equivalent number of CFUs from WT, *iniA∷hyg* and complemented strains grown in LIMM. ST142 growth was determined based on the number of viable cells detected by the resazurine cell viability assay. (**B**). Growth of WT (filled symbols) and *iniA∷hyg* (open symbols) in high Fe (circles) and low Fe (squares) MM based on increase in optical density (O.D). Data represents the mean ± SD of three independent cultures. (**C**). Levels of mycobactin extracted from WT and *iniA∷hyg* grown in low iron conditions. Shown is the average of mycobactin extracted from two independent cultures. \*\*\*\**P* < 0.00005.

### Reduced release of bacteria-derived EV by *iniA∷hyg* infected macrophages

Previous studies have shown that bacteria-derived EV are released by *Mtb*-infected mouse macrophages and can act as potent immunomodulators of uninfected cells (5, 6). To determine whether *Mtb* cells require DLPs to produce EV in macrophages, we compared the abundance of vesicles containing bacterial components released into the culture medium from THP-1 macrophages 24 hours after infection with different *Mtb* strains: WT, *iniA∷hyg*, or *iniA∷hyg* complemented with *iniBAC. Mtb* derived EV were less abundant in the medium of *iniA∷hyg*-infected macrophages compared to macrophages infected with WT or *iniBAC*-complemented *iniA∷hyg Mtb* (Fig. 8*B*). This effect was not due to reduced replication of the mutant, as WT and *iniA∷hyg* exhibit equivalent growth in THP-1 cells (Fig. 8*A*). These findings indicate DLPs are not required for intracellular survival and replication of *Mtb* but are necessary for production of bacteria-derived vesicles released by *Mtb*-infected macrophages.

**Figure 8.**
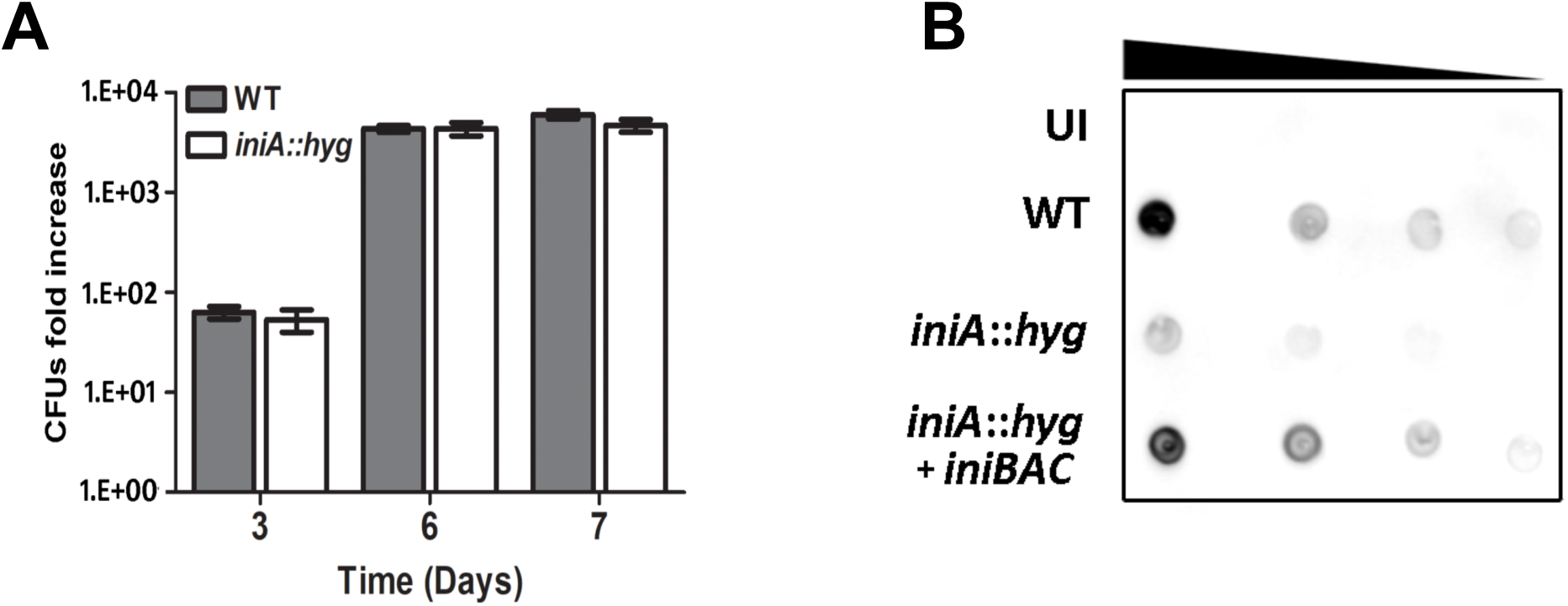
Growth and EV production during macrophage infection. (**A**). Increase in CFUs at the indicated times relative to time zero of WT and *iniA∷hyg* infecting THP-1 cells. (**B**). Detection of *Mtb*-EV in the culture medium of uninfected (UI) or THP-1 infected with WT, *iniA:hyg* and complemented *Mtb* by dot-blot, using an anti-EV polyclonal murine serum. Black triangle indicates increasing dilutions of culture supernatant loaded in the membrane. Data are representative of two independent experiments performed in triplicate.

## DISCUSSION

There is compelling evidence suggesting that EV produced by pathogenic mycobacteria constitute a delivery mechanism for immunologically active molecules and bacterial factors that contribute to virulence. However, we have very little understanding of the mechanisms and regulation of EV biogenesis in mycobacteria. By analyzing cellular features shared between *Mtb* cells under conditions that stimulate EV production, we uncovered changes in cell envelope ultrastructure associated with vesicle production and identified DLPs as essential factors in EV biogenesis in culture and during infection of macrophages.

Proteins of the dynamin superfamily mediate membrane fusion, fission and membrane restructuring. Dynamin, the founding member of this family, mediates membrane fission leading to the budding of vesicles during endocytosis (29). Bacterial DLPs are poorly understood, but they appear to function in a variety of processes: *Escherichia coli* CrfC functions in chromosome partitioning during DNA replication (30), DynA and DynB in *Streptomyces* are involved in cell division during sporulation (31), DynA in *Bacillus subtillis* can tether membranes and may mediate membrane repair (32), and LeoA, B and C in enterogenic *E. coli* localize to the periplasm and contribute to virulence through membrane vesicle associated toxin secretion (33).

Early studies showed *Mtb*-IniA assemble into hexameric-ring like structures upon interaction with lipid monolayers (23) and recent structural and biochemical analysis of *Msm*-IniA confirmed protein folding and biochemical characteristics of DLPs including GTPase activity, interaction with membrane lipids and GTP-hydrolysis dependent membrane fission activity *in vitro* (22). Most bacterial DLPs exist as side by side pairs within operons like *iniAC* or in the case of *B. subtillis* DnyA, as two DLPs genetically fused into a single unit (21) which suggest interactions of DLP isoforms for membrane remodeling. The significance of DLP association was demonstrated for the *Campilobacter jejuni* DLPs. Both *Cj*-DLPs are monomeric in solution and form a stable tetramer when mixed. This tetramer tethers and bridges distantly opposing membranes (34). Our complementation analyses indicated IniA and IniC have non-redundant functions in EV biogenesis supporting the idea that these DLPs work together. This is in agreement with Wang et al. who observed low GTPase activity in monomeric *Msm*-IniA and suggested that interactions with IniC may be required for high rates of GTP hydrolysis (22). This form of cooperation was described for mitochondrial OPA1 proteins. A long and a short isoform corresponding to membrane-anchored and soluble OPA1 form a complex and while the long isoform provides membrane anchoring, the short isoform contributes a strong GTPase activity for mitochondrial inner membrane fusion (35). These observations and our findings, suggest a model in which like dynamin, IniA polymerizes on the membrane bilayer and working with IniC as a mechanochemical GTPase complex mediates the membrane fission necessary for release of mycobacterial EV. The interactions between IniA and IniC and the mechanistic details underlying DLP function in EV release are currently under investigation.

Our ultrastructural analysis of hypervesiculating *Mtb* cells suggested a close link between EV release and cell envelope alterations. Our microscopy observations that indicate loss of OM thickness in Fe-limited *Mtb* are consistent with previous analysis of the cell envelope of Fe deficient *Mtb* which showed reduced cell envelope lipids (16) and enhanced cell envelope permeability (16, 36) associated with downregulation of *mmpL3* (17). MmmpL3 transports trehalose monomycolate across the plasma membrane for outer membrane assembly (37). Our transcriptomic analysis showed additional genes required for MA and glycolipid biosynthesis that are repressed in Fe-limited *Mtb* (Fig S1) (16, 17). Modification of cell envelope lipids during Fe-limitation may directly influence EV release by increasing cell envelope permeability and through induction of *iniBAC*. It has been shown that *iniBAC* induction is mediated by activation of IniR (***ini****BAC* Regulator), a signal transduction ATPase with a sugar-binding domain (38). It was postulated that inhibition of MA biosynthesis leads to reduction in trehalose dimycolate, and consequently increased levels of free trehalose, which can induce *iniBAC* expression via IniR activation (38). Induction of *iniBAC* during iron limitation is compatible with this model, as repression of MA and TDM synthesis might lead to the accumulation of free trehalose.

The ultrastructural alterations in the *virR* mutant combined with the changes in PG integrity demonstrated here, and the reported function of other LCP family members, all point to changes in PG dynamics in connection with enhanced vesicle release. Thus, collectively our findings in Fe-limited and VirR-restricted *Mtb* cells suggest enhanced EV release in association with outer membrane or cell wall alterations. In this regard, enhanced EV release has also been reported in *Mtb* under conditions that activate the phosphate-sensing, two-component system SenX3-RegX3 (39). Hypersensitivity to surface stress has been observed associated with activation of RegX3, suggesting this system is also linked to cell envelope integrity (40). It would be interesting to examine cell envelope ultrastructural changes in the context of RegX3 activation.

The association of EV production and cell envelope restructuring is evidenced also during exposure to the antibiotic INH. Subinhibitory concentrations of this MA synthesis inhibitor stimulated EV release and this effect was dependent on induction of *iniBAC*. This observation may have relevant implications for TB treatment as antibiotic-stimulated production of EV and their proinflammatory effects may undermine immune control and compromise antibiotic effectiveness. Overexpression of *iniA* moderately increased *Msm* and BCG tolerance to INH and under specific conditions the *Mtb iniA∷hyg* mutant showed increased sensitivity to INH (23). Whether the effect of IniA in regulating INH sensitivity is linked to its function as DLP merits further investigation.

Finally, our data showing the release of vesicles containing bacterial components into the medium by *Mtb*-infected macrophages and that vesicle-mediated Fe-acquisition is dependent on DLPs, highlights the potential relevance of DLPs during infection. Accordingly, IniA and IniC were found induced in the lungs of infected guinea pigs (41). Since bacteria-derived vesicles released by *Mtb*-infected macrophages have been previously demonstrated to regulate uninfected macrophages and inhibit T cell activation (5, 6), our findings suggest that targeting DLPs-mediated vesicle production might effectively impede the immunomodulatory ability of *Mtb* and thereby, may enhance immune control of TB infection.

## MATERIALS AND METHODS

### Bacterial Strains, media and growth conditions

*Mtb* H37Rv (ATCC) and derivative strains *virR∷Tn* (9) and *iniA∷hyg* (23) were used in this study. Iron (Fe)-depleted minimal medium (MM) was used for culture under iron-controlled conditions. MM contains 0.5% (wt/vol) asparagine, 0.5% (wt/vol) KH_2_PO_4_, 0.1% glycerol, 0.05% Tyloxapol, 0.2% (wt/vol) dextrose, 0.085% NaCl, pH to 6.8. To remove metal ions MM was treated with 5% Chelex-100 (BioRad) for 24hr, at 4°C with gentle agitation. Chelex-100 was removed by filtration through 0.22 µm filter (Millipore) and the medium was supplemented with sterile 0.5 mg of ZnCl_2_ liter ^-1^, 0.1 mg of MnSO_4_ liter^-1^, and 40mg of MgSO_4_ liter^-1^. The amount of residual Fe in this medium, determined by atomic absorption spectroscopy is ∼1 µM. When indicated high Fe MM was prepared by supplementing Fe-depleted MM with 50 µM FeCl_3_.

### Complementation of *iniA*∷*hyg Mtb*

The *Mtb iniBAC* operon and its native promoter (included in 211 bp upstream *iniB*) was PCR amplified from chromosomal DNA using primers *iniBAC*_*TB*_ *com F* and *iniBAC*_*TB*_ *com R* (Table S1). The PCR product was cloned at the StuI site of pSM316 (42), a vector that integrates at the *attB* site. The resulting plasmid, pSM986, was electroporated into *iniA*∷*hyg*. Transformants were selected on 7H10 agar with streptomycin (20 µg.ml^-1^) and spectinomycin (75 µg.ml^-1^). *iniA* and *iniC* complementing plasmids were generated by PCR amplification of each gene and cloning into pMV361 in front of the hsp60 promoter using Fast cloning (43). The resulting plasmids pSM988 and pSM989 were electroporated into *iniA*∷*hyg* and transformants selected on 7H10 agar containing 20 µg.ml^-1^ of kanamycin. All plasmids were validated by DNA sequencing before electroporation.

### Mycobactin determination

Mycobacterial strains were grown to mid-logarithmic phase in 7H9 medium, and 0.7 ml of culture was spread on MM agar. After incubation at 37° for 10 days, the bacteria were scraped from the plate. Subsequently, mycobactin was extracted in ethanol and chloroform and quantified as previously described and normalized to the weight of the bacterial pellet (44).

### Electron-Microscopy

Cells were fixed with 2% glutaraldehyde in 0.1 M cacodylate at room temperature for 24 h, and then incubated overnight in 4% formaldehyde, 1% glutaraldehyde, and 0.1% PBS. For scanning microscopy, samples were then dehydrated through a graded series of ethanol solutions before critical-point drying using liquid carbon dioxide in a Toumisis Samdri 795 device and sputter-coating with gold-palladium in a Denton Vacuum Desk-2 device. Samples were examined in a Zeiss Supra Field Emission Scanning Electron Microscope (Carl Zeiss Microscopy, LLC North America), using an accelerating voltage of 5 KV.

For Cryo-EM, grids were prepared following standard procedures and observed at liquid nitrogen temperatures in a JEM-2200FS/CR transmission electron microscope (JEOL Europe, Croissy-sur-Seine, France) operated at 200 kV. An in-column omega energy filter helped to record images with improved signal/noise ratio by zero-loss filtering. The energy selecting slit width was set at 9 eV. Digital images were recorded on an UltraScan4000 CCD camera under low-dose conditions at a magnification of 55,058 obtaining a final pixel size of 2.7 Å/pixel.

Density profiles were calculated along rectangular selections with the ImageJ software (NIH, Bethesda, MD). Average and standard error for each strain was calculated on measurements from 50 independent cells.

### RNA sequencing

*Mtb* strains were grown in MM supplemented with 0.05% Tyloxapol at 37°C until they reached an O.D. 595nm of 0.3 and harvested by centrifugation. The cell pellets were resuspended in 1 ml Qiagen RNA protect reagent (Qiagen) and incubated for 24 h at room temperature. Cells were disrupted by mechanical lysis in a FastPrep-24 instrument (MP Biomedicals, Santa Ana, CA) in Lysing Matrix B tubes and RNA was purified with the Direct-zol RNA miniprep kit (Zymo Research, Irvine, CA). The quantity and quality of the RNAs were evaluated using Qubit RNA HS Assay Kit (Thermo Fisher Scientific) and Agilent RNA 6000 Nano Chips (Agilent Technologies), respectively. Sequencing libraries were prepared using the Ribo-Zero rRNA Removal Kit (Gram-positive Bacteria) (Illumina Inc.) and the TruSeqStranded mRNA library prep kit (Illumina Inc), following the Ribo-Zero rRNA Removal kit Reference guide and the “TruSeqStranded mRNA Sample Preparation Guide. Briefly, starting from 1 µg of total RNA, bacterial rRNA was removed and the remaining RNA was cleaned up using AgencourtRNAClean XP beads (Beckman Coulter). Purified RNA was fragmented and primed for cDNA synthesis. cDNA first strand was synthesized with SuperScript-II Reverse Transcriptase (Thermo Fisher Scientific) for 10 min at 25°C, 15 min at 42°C, 15 min at 70°C and pause at 4°C. cDNA second strand was synthesized with Illumina reagents at 16°C for 1 hr. Then, A-tailing and adaptor ligation were performed. Libraries enrichment was achieved by PCR (30 sec at 98°C; 15 cycles of 10 sec at 98°C, 30 sec at 60°C, 30 sec at 72°C; 5 min at 72°C and pause at 4°C). Afterwards, libraries were visualized on an Agilent 2100 Bioanalyzer using Agilent High Sensitivity DNA kit (Agilent Technologies) and quantified using Qubit dsDNA HS DNA Kit (Thermo Fisher Scientific). Library sequencing was carried out on an Illumina HiSeq2500 sequencer with 50 nucleotides single end reads and at least 20 million of reads per individual library were obtained.

### RNA-sequencing data analysis

Quality Control of sequenced samples was performed by *FASTQC* software (http://www.bioinformatics.babraham.ac.uk/projects/fastqc/). Reads were mapped against the *M. tuberculosis* H37Rv strain (*GCF_000195955*.*2_ASM19595v2)* reference genome using Tophat (Trapnell, 2009) with --bowtie1 option, to align 50 bp reads. The resulting *BAM* alignment files for the samples were the input to Rsubread’s (45) featureCounts function to generate a table of raw counts required for the Differential Expression (DE) analysis, carried out by DESeq2 (46), to detect differentially expressed genes (FDR< 0.05 and Foldchange> 1.5) among the different conditions.

### Real-Time RT-PCR analysis

RNA was obtained as above and reverse-transcribed using M-MLV reverse transcriptase (Thermo Fisher Scientific). Real-time PCR was then performed using SYBR Green PCR Master Mix (Thermo Fisher Scientific) on a QuantStudio 6 real-time PCR System (Thermo Fisher Scientific). Fold induction of *iniA* was calculated using the 2^−ΔΔCt^ method (47) relative to the reference gene, P1-*rrnA* in the case of *iniA* expression in cells treated with INH and relative to sigA in RT-PCR that validated RNA-seq gene expression. Primers were designed using the Primer3 software (48).

### Muropeptides release assay

The method described by Hadzija (49) was applied with some modifications. Mycobacteria were cultured in 10 ml of 7H9 medium. When cultures reached an O.D of 0.5, cells were washed in MM, expanded in 100 ml of high Fe MM and incubated at 37°C for 5 additional days. Then, bacterial cells were harvested and plated to determine CFUs. Cultured supernatants were filtered through 0.22 µm and lyophilized. Lyophilized material was resuspended in 1 ml of mQ water, mixed with 5 ml of H_2_SO_4_ and boiled for 30 min. After cooling, 0.05 ml of 4% (w/v) CuSO4 in H_2_O and 0.1 ml of 1.5% (w/v) p-hydroxydiphenyl in ethanol were added and incubated at 30°C for 30 min. Absorbance was measured at 560 nm and normalized to O.D and CFUs.

### Lysozyme susceptibility assay

Mtb WT and *virR*∷*tn* were grown in 7H9 media supplemented with ADC and tween 80 to an O.D 540 of 0.2 and then diluted to a O.D 540 of 0.1 with 7H9 containing 0 or 50 µg.ml^-1^ lysozyme (Sigma). The cultures were incubated with agitation at 37°C and O.D 540 was monitored daily.

### Comparative Modeling

The *Mtb*-IniA (36-628) comparative model was generated using Phyre2 (50). The *Msm*-IniA X-ray crystal structure (Protein data bank code 6J73) was used as the modeling template. The *Mtb*-IniC (1-493) comparative model was generated using Phyre2 (50) using the *Msm*-IniA X-ray crystal structure (Protein data bank code 6J73) as the modeling template. Molecular graphics were produced with Pymol 2.1.

### EV isolation and purification

*Mtb*-EV were prepared and purified as previously (51). Briefly, the culture filtrate of 1 L cultures of *Mtb* grown in low iron MM without detergent for 14 days was processed for vesicle isolation by differential centrifugation. In parallel a 2 ml culture in MM supplemented with 0.05% tyloxapol to disperse bacterial clumps was used to determine viability by plating culture dilutions onto 7H10 agar plates and enumerating colony forming unit (CFU) at the time of culture filtrate collection.

The membranous pellet containing vesicles obtained after ultracentrifugation of the culture filtrate at 100,000 ×*g* for 2 h at 4°C, was resuspended in 1 ml sterile phosphate-buffered saline (PBS) and overlaid with a series of Optiprep gradient layers with concentrations ranging from 45% to 20% (w/v). The gradients were centrifuged at 100,000 × g for 16 h. At the end 1 ml fractions were removed from the top, diluted to 20 ml with PBS and purified vesicles recovered by sedimentation at 100,000 × g for 1 h. Vesicle pellets were suspended in PBS before analysis. Protein concentration was measured by Bradford assay (Bio Rad) according to manufacturer instructions.

### EV-lipid determination

The lipophilic fluorescent membrane probe 1-(4-Trimethylammoniumphenyl)-6-Phenyl-1,3,5-Hexatriene *p*-Toluenesulfonate (TMA-DPH) (Themo Fisher Scientific) was used to assess EV associated lipids. 10 µl of vesicle preparations and TMA-DPH at a final concentration of 50 µM in 100 µl of PBS were mixed in 96 well black plates and incubated at 33°C for 20 min before measuring fluorescence in a Synergy HI (Biotek) plate reader at 360 nm excitation-430 nm emission.

### Nano particle tracking analysis

Particle concentration and size of Mtb-EV preparations were measured using ZetaView (Particle Metrix) at 23.85 °C and pH 7,0. Before the measurement, standard silica beads (100 nm) were measured and the instrument was calibrated. Samples were diluted in pre-filtered PBS to approximately 10^6^-10^7^ particles.ml^-1^ in Millipore DI water. Triplicate videos of each sample were taken in light scatter mode. Particle size and concentrations were analyzed using the built-in EMV protocol and plotted using graph pad prism 5.0 software.

### Western Dot blot

2 µl of EV isolates and two fold serial dilutions were spotted onto a nitrocellulose membrane (Abcam) and process for dot blot using anti-EV polyclonal murine serum (8) as primary antibody and goat anti-mouse-HRP conjugated as secondary antibody and ECL prime Western Blotting Detection chemiluminescent substrate (GE Healthcare). The signal was visualized in a Chemidoc MP imaging system (BIO-RAD).

### EV mediated Fe-transfer assay

The siderophore mutant (ST142) was pre-incubated in LIMM supplemented with 100 µg.ml^-1^ Hygromycin B for 24 h. Approximately, 5 ×10^4^ ST142 cells in a volume of 100 µl were seeded in 96 well plates in triplicate and incubated with EV isolated from the culture filtrate of 1 × 10^9^ CFUs of WT, *iniA*∷*hyg* or complemented strains at 37°C for 5 days. Bacterial growth was assessed at day five using the resazurin assay (52).

### Stimulation of EV release by isoniazid

*Mtb* H37Rv was grown in high iron MM with 0.05% Tyloxapol. When cultures reached an O.D. 540 of 0.5, aliquots of 1 ml were washed in PBS with 0.05% Tyloxapol and resuspended in 1 ml of fresh MM without detergent, supplemented with various concentrations of isoniazid (INH) and incubated at 37°C for 5 days. For each concentration of antibiotic cell viability was assessed by the resazurin assay (52). EV release was determined by western dot-blot and vesicle associated lipid and protein was determined in vesicles isolated from a 50 ml culture containing 50 ng.ml^-1^ of INH

### THP-1 infection and *Mtb*-EV detection

Infection of THP-1 human monocytic cells was performed as previously described (53). Briefly, 1×10^5^ THP-1 cells per well were differentiated with 50 nM phorbol myristate acetate in 96 well plates and infected with *Mtb* strains, pre-grown in 7H9 medium to logarithmic phase, at a multiplicity of infection of 0.05 bacterium per macrophage. After 4 hours of incubation at 37°C in 5% CO_2_ atmosphere, the medium was removed, and the cells were extensively washed with pre-warmed PBS to remove extracellular bacteria. At indicated time points after infection, triplicate wells for each *Mtb* strain infection were treated with 0.05% sodium dodecyl sulfate (SDS) to lyse the macrophages and numbers of CFU were determined by plating serial dilutions on 7H10 plates. For *Mtb*-EV detection 2×10^7^ THP-1 macrophages were infected with *Mtb* strains at MOI of 2. 24hr post infection the culture medium was collected filtered sterilized and processed for EV isolation by differential centrifugation. Preparations of EV were analyzed by semiquantitative dot blot after normalization to macrophage total protein and bacterial CFUs 24h post infection.

## Statistical analysis

The statistical significance of the difference between experimental groups was determined by the two-tailed Student’s test using PRISM 5.0. *P* values *≤* than 0.05 were considered significant. Statistical analysis of RNAseq is detailed in the corresponding section of Materials and Methods.

## Data Availability

All data presented is available. Data from RNAseq will be deposited at Gene expression Omnibus (GEO).

## Founding Information

This work was supported by a National Institute of Health Grant R21 130628 (G. M.R). R.P.-R. is further a ‘Ramon y Cajal’ fellow from the Spanish Ministry of Economy and Competitiveness. R.P.-R. acknowledges support by MINECO/FEDER EU contracts SAF2016–77433-R. CIC bioGUNE thanks MINECO for the Severo Ochoa Excellence Accreditation (SEV-2016–0644). JA: SAF2015-65327-R and RTI2018-096494-B-100 from MINECO. AP holds a fellowship form the Department of Education of the Basque Government (PRE_2018_1_0229).

## Acknowledgments

We are very grateful to Carl Nathan, David Alland and Roberto Colangeli (Rutgers University) for sharing mutant *Mtb* strains and David Dubnau for helpful discussions and review of the manuscript.

The authors have no conflict of interest to declare.

## SUPPLEMENTARY FIGURE LEGENDS

**Supplementary Figure S1. Repression of cell envelope lipid biosynthesis genes in response to Fe limitation**. Shown is the fold change in the abundance of transcripts corresponding to the indicated genes at day 1 and 7 during Fe deprivation relative to cells maintained in Fe sufficient conditions. The figure was crated based on reanalysis of published data (15).

**Supplementary Figure S2. Sensitivity to lysozyme and muropeptide release in *virR***. (**A**) Growth of WT and *virR* treated with 0 or 50 µl.mL^-1^ lysozyme (lys). (**B**) Growth (top panel) and release of PG derived muramic acid into the extracellular medium (lower panel) of WT (blue circles), *virR* mutant (orange circles) and complemented strain (open circles). Data represents the mean ± SD of three independent cultures.

**Supplementary Figure S3. Comparative gene expression in *virR* and LI *Mtb***. The pool of genes upregulated or downregulated in *virR* compared to differentially expressed genes in *Mtb* grown under low versus high iron conditions for 24 h (**A**) and 1 week (**B**) based on the trascriptomic analysis of *virR* carried out here and data of iron deprived *Mtb* published in (15). Color code for both set of data indicate significant upregulation (blue) or downregulation (red). Transcriptomic datasets were log2 transformed and conditionally formatted to provide the color scale based on maximum and minimum values.

**Supplementary Figure S4. Growth of *virR* in Fe limiting conditions**. Growth of *virR* mutant and WT in low iron MM monitored by increase in O.D 540 nm. Data represents the mean ± SD of three independent cultures.

**Supplementary Figure S5. EV purification from WT, *iniA∷hyg* and complemented strains**. (**A**) Photograph of crude EV pellets prepared from the culture filtrate of WT, *iniA∷hyg* and *iniBAC* complemented strains normalized to CFUs, showing drastic reduction in *iniA∷hyg*. (**B**) Protein and (**C**) lipid content in each density gradient fraction after separation of crude EV preparations from WT, *iniA∷hyg* and complemented strains. (**D**) Nanoparticle analysis of density fraction 3 of from WT and *iniA∷hyg*. (E-G) Protein profile in each density gradient fraction separated by SDS-PAGE and stained with Sypro Ruby from indicated strains showing below each gel a dot-blot of each density gradient fraction using an anti-EV polyclonal murine serum as described in materials and methods.

**Supplementary Figure S6. Secreted protein in the culture supernatant of *Mtb* strains**.

Protein in the culture supernatant and in the EV fraction from WT and *iniA*∷hyg. Data represents the mean ± SD of three independent cultures.

**Supplementary Figure S7. INH dependent *iniA* induction**. Induction of *iniA* in *Mtb* cells treated with increasing concentrations of INH in Fig. 5B, determined by RT-PCR. Fold change was calculated relative to untreated cells.

