## Supplemental Figures for "Dynamin-like proteins are essential for vesicle biogenesis in *Mycobacterium tuberculosis*"

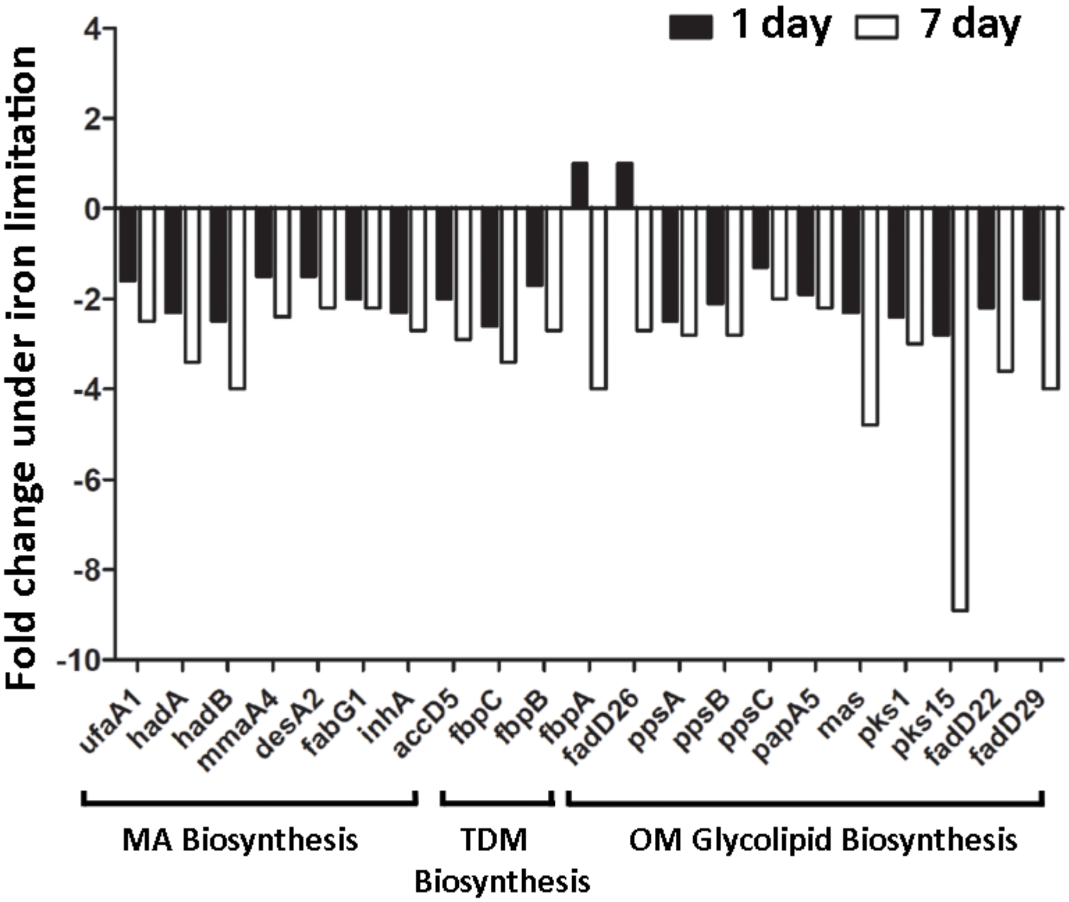


**Supplementary Figure 1. Repression of cell envelope lipid biosynthesis genes in response to Fe limitation.** Shown is the fold change in the abundance of transcripts corresponding to the indicated genes at day 1 and 7 during Fe deprivation compared to cells maintained in Fe sufficient conditions. The figure was crated based on reanalysis of data published previously (15).

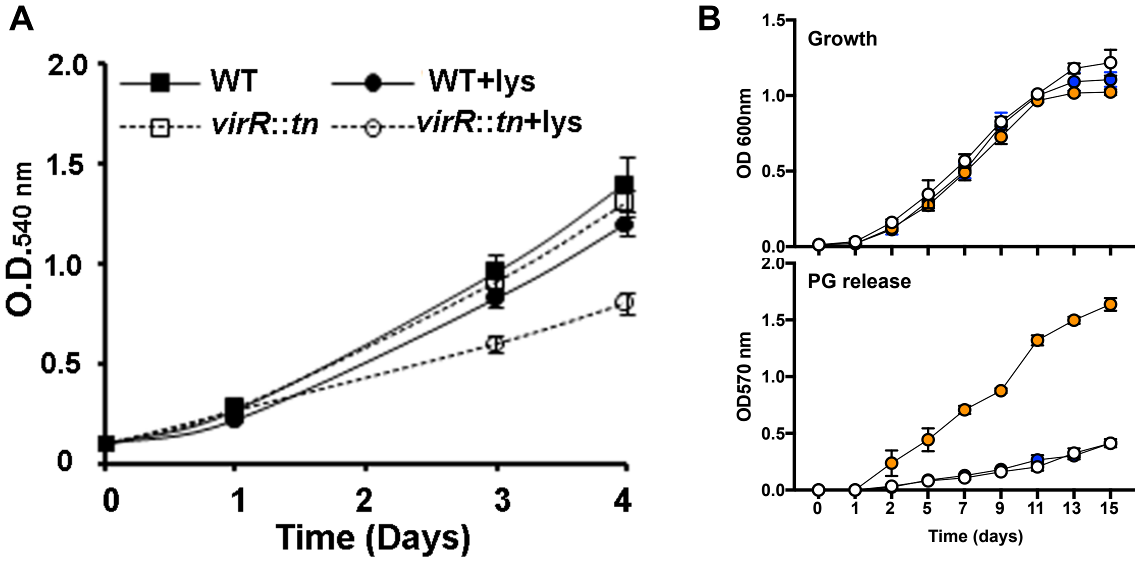


**Supplementary Figure 2**. **Sensitivity to lysozyme and muropeptide release in *virR*.** (**A**) Growth of WT and *virR* treated with 0 or 50 µl.mL$^{-1}$ lysozyme (lys). (**B**) Growth (top panel) and release of PG derived muramic acid into the extracellular medium (lower panel) of WT (blue), *virR* mutant (orange) and complemented strain (open). Data represents the mean ± SD of three independent cultures.

**A**

| GENE_SYMBOL | virR mut | Low Fe 24h | |
|---|---|---|---|
| Rv0079 | -1,105 | 1,617 | |
| Rv0080 | -1,676 | 0,932 | |
| Rv0081 | -1,254 | 0,301 | |
| oxcA | 0,723 | 1,675 | |
| bglS | -0,919 | -0,537 | |
| PPE3 | -0,774 | 0,739 | |
| Rv0326 | -1,136 | 1,241 | |
| Rv0331 | -1,158 | 0,238 | |
| iniB | 1,123 | 0,307 | *iniBAC* |
| iniC | 0,677 | 0,304 | |
| grpE | -0,813 | -0,356 | |
| Rv0431 | 0,080 | -1,678 | *virR* |
| sodC | 1,226 | -0,495 | |
| Rv0574c | -1,151 | 0,318 | |
| Rv0601c | 0,681 | 0,639 | |
| fadE9 | -0,899 | 1,339 | |
| PE_PGRS11 | -0,798 | 0,417 | |
| Rv0763c | -0,731 | 0,218 | |
| cyp51 | -0,833 | -1,255 | |
| Rv0765c | -0,996 | 0,237 | |
| cyp123 | -0,920 | 0,091 | |
| aldA | -1,214 | 0,341 | |
| Rv0790c | -0,974 | 0,464 | |
| Rv0791c | -1,091 | 0,382 | |
| Rv0792c | -1,206 | 0,632 | |
| Rv0996 | -0,662 | -0,669 | |
| prpD | 1,059 | -1,691 | |
| prpC | 0,809 | -1,769 | |
| PPE17 | -0,916 | 0,954 | |
| lipX | -1,252 | 1,097 | |
| sigE | -0,968 | -0,466 | |
| cysD | -1,140 | 1,199 | |
| Rv1286 | -0,953 | 1,122 | |
| Rv1303 | -0,676 | -0,525 | |
| PE_PGRS27 | 0,682 | 0,299 | |
| Rv1460 | 0,954 | 2,463 | |
| Rv1462 | 0,738 | 2,596 | *S-Fe cluster* |
| Rv1463 | 0,779 | 2,719 | |
| csd | 0,771 | 2,212 | |
| Rv1465 | 0,682 | 2,569 | |
| Rv1466 | 0,659 | 2,450 | |
| Rv1577c | -0,691 | 0,500 | |
| Rv1586c | -0,817 | -0,162 | |
| cydC | -1,156 | 0,814 | |
| Rv1733c | -1,365 | 0,110 | |
| Rv1734c | -2,042 | 0,085 | |
| Rv1738 | -1,328 | 1,862 | |
| PE20 | -0,731 | 1,403 | |
| Rv1856c | 0,876 | 0,790 | |
| Rv1954c | -1,253 | 0,505 | |
| Rv1996 | -2,154 | 0,661 | |
| Rv1999c | -0,727 | 0,169 | |
| Rv2016 | -0,949 | 0,613 | |
| Rv2017 | -0,782 | 0,707 | |
| hspX | -0,981 | 1,382 | |
| sigC | -0,691 | -0,598 | |
| ftsQ | -0,754 | -0,491 | |
| Rv2516c | -0,712 | 0,916 | |
| Rv2517c | -0,879 | 1,314 | |
| TB31.7 | -1,398 | 1,144 | |
| Rv2624c | -1,573 | 0,115 | |
| Rv2625c | -1,343 | 0,099 | |
| hrp1 | -1,221 | 0,780 | |
| Rv2627c | -1,533 | 0,425 | |
| Rv2631 | -1,011 | 0,062 | |
| Rv2651c | -0,683 | 0,544 | |
| Rv2660c | -0,904 | 0,930 | |
| dacB2 | -0,800 | -0,626 | |
| Rv2912c | -1,223 | -0,450 | |
| ppsB | 0,787 | -1,266 | |
| ppsC | 0,977 | -1,340 | |
| ppsE | 0,716 | 0,106 | |
| drrA | 0,701 | -2,675 | |
| Rv2990c | -0,985 | 0,407 | |
| PPE48 | 0,861 | 0,367 | |
| Rv3128c | -1,059 | 0,869 | |
| tgs1 | -1,385 | 1,377 | |
| Rv3131 | -1,193 | 1,250 | |
| Rv3137 | 0,678 | -1,042 | |
| Rv3169 | -0,715 | 0,495 | |
| rubA | -0,796 | 1,473 | |
| alkB | -0,811 | 2,393 | |
| ctpC | -0,680 | -0,498 | |
| usfY | -0,960 | 1,481 | |
| lat | -1,163 | 3,322 | |

**B**

| GENE_SYMBOL | virR mut | Low Fe 1 w | |
|---|---|---|---|
| Rv0325 | -1,120 | 0,779 | |
| Rv0331 | -1,158 | 0,819 | |
| Rv0340 | 0,739 | 2,286 | |
| iniA | 0,966 | 1,720 | *iniBAC* |
| iniC | 0,677 | 0,867 | |
| Rv0571c | -0,981 | 1,018 | |
| Rv0574c | -1,151 | 0,712 | |
| mce2R | -0,967 | 1,040 | |
| fadE9 | -0,899 | 0,876 | |
| mmsA | -1,048 | 1,398 | |
| PE_PGRS11 | -0,798 | 1,002 | |
| Rv0763c | -0,731 | 0,949 | |
| Rv0765c | -0,996 | 0,797 | |
| aldA | -1,214 | 0,611 | |
| Rv0791c | -1,091 | 0,729 | |
| Rv0793 | -1,437 | 0,762 | |
| prpC | 0,809 | -1,301 | |
| PPE17 | -0,916 | 0,864 | |
| lipX | -1,252 | 0,955 | |
| sigE | -0,968 | 0,622 | |
| cysD | -1,140 | 2,534 | |
| PE_PGRS27 | 0,682 | 1,016 | |
| Rv1460 | 0,954 | 4,599 | |
| Rv1462 | 0,738 | 2,364 | |
| Rv1463 | 0,779 | 2,666 | *S-Fe cluster* |
| csd | 0,771 | 3,088 | |
| Rv1465 | 0,682 | 3,041 | |
| Rv1466 | 0,659 | 3,658 | |
| Rv1577c | -0,691 | 1,627 | |
| Rv1585c | -0,723 | 1,575 | |
| Rv1586c | -0,817 | 1,026 | |
| cydC | -1,156 | 0,757 | |
| Rv1733c | -1,365 | 0,750 | |
| PE20 | -0,731 | 3,220 | |
| higB | -1,140 | 1,589 | |
| higA | -1,154 | 1,495 | |
| ctpG | -1,572 | 1,551 | |
| Rv1993c | -1,633 | 3,671 | |
| cmtR | -1,912 | 4,611 | |
| Rv2017 | -0,782 | 0,934 | |
| ftsQ | -0,754 | -0,312 | |
| Rv2516c | -0,712 | 2,242 | |
| Rv2517c | -0,879 | 3,912 | |
| Rv2651c | -0,683 | 2,485 | |
| Rv2660c | -0,904 | 4,482 | |
| Rv2913c | -1,419 | 1,843 | |
| fadE22 | -1,151 | 0,838 | |
| Rv3188 | 1,102 | 1,961 | |
| usfY | -0,960 | 2,628 | |
| Rv3289c | -1,197 | 3,027 | |
| lat | -1,163 | 4,885 | |
| Rv3833 | -0,937 | 1,065 | |

**Supplementary Figure 3**. **Comparative gene expression in *virR* and LI Mtb.** The pool of genes upregulated or downregulated in virR compared to differentially expressed genes in Mtb grown under low versus high iron conditions for 24 h (**A**) and 1 week (**B**) based on the trascriptomic analysis of virR carried out here and data of iron deprived Mtb published in (15). Color code for both set of data indicate significant upregulation (blue) or downregulation (red). Transcriptomic datasets were log2 transformed and conditionally formatted to provide the color scale based on maximum and minimum values.

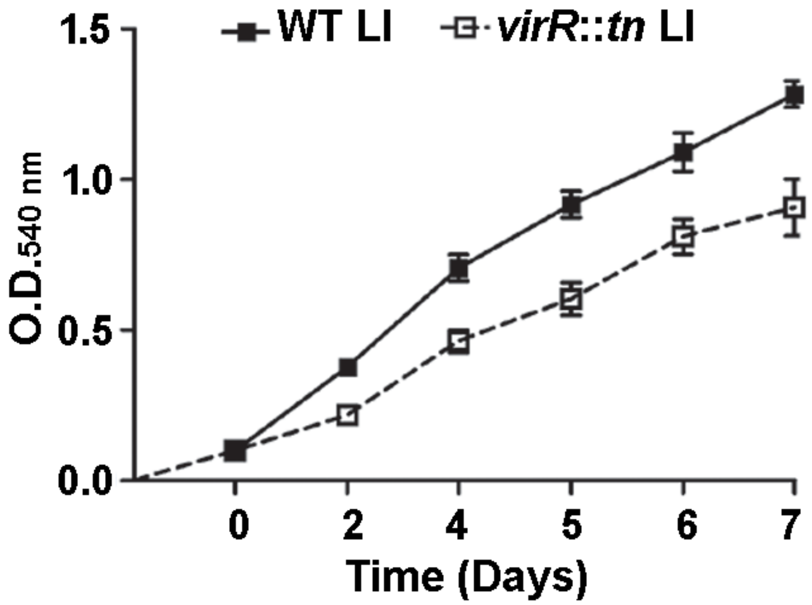


**Supplementary Figure 4. Growth of *virR* in Fe limiting conditions.** Growth of *virR* mutant and WT in low iron MM monitored by increase in O.D 540 nm. Data represents the mean ± SD of three independent cultures.

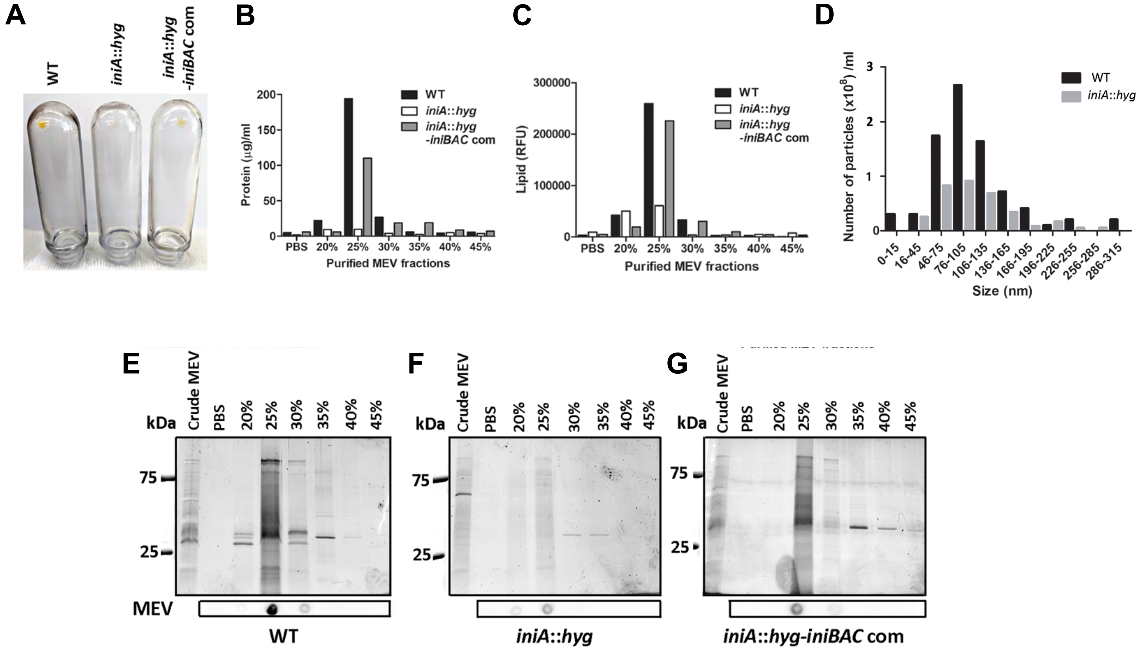


**Supplementary Figure 5. EVs purification from WT, *iniA::hyg* and complemented strains.** (**A**) Photograph of crude membrane vesicle pellets prepared from the culture filtrate of WT, *iniA::hyg* and *iniBAC* complemented strains normalized to CFUs, showing drastic reduction in *iniA::hyg*. (**B**) Protein and (**C**) lipid content in each density gradient fraction after separation of crude membrane vesicle preparations from WT, *iniA::hyg* and complemented strains. (**D**) Nanoparticle analysis of density fraction 3 of from WT and *iniA::hyg*. (E-G) Protein profile in each density gradient fraction showing below each gel a dot-blot of each density gradient fraction using an anti-EVs polyclonal murine serum as described in materials and methods.

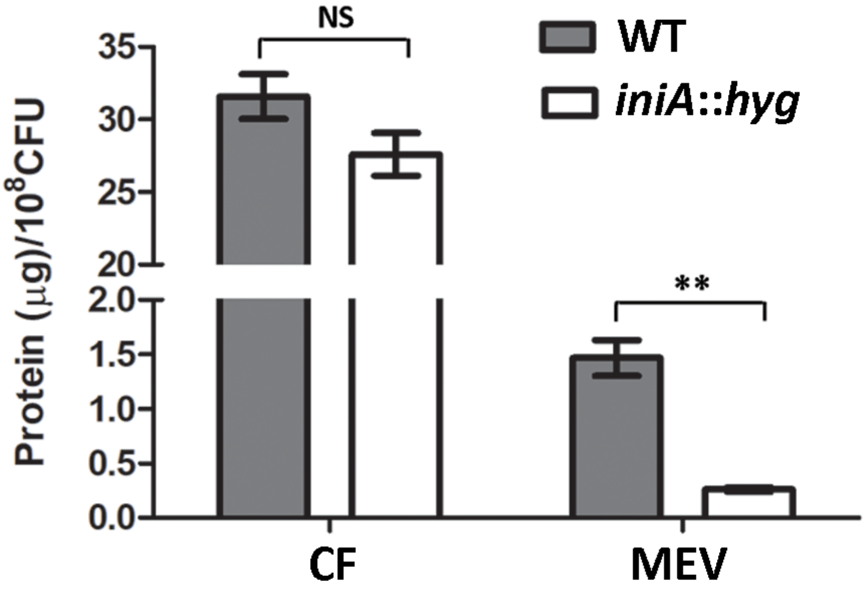


**Supplementary Figure 6. Secreted protein in the culture supernatant of *Mtb* strains.** Protein in the culture supernatant and in the MEVs fraction from WT and *iniA*::hyg. Data represents the mean ± SD of three independent cultures.

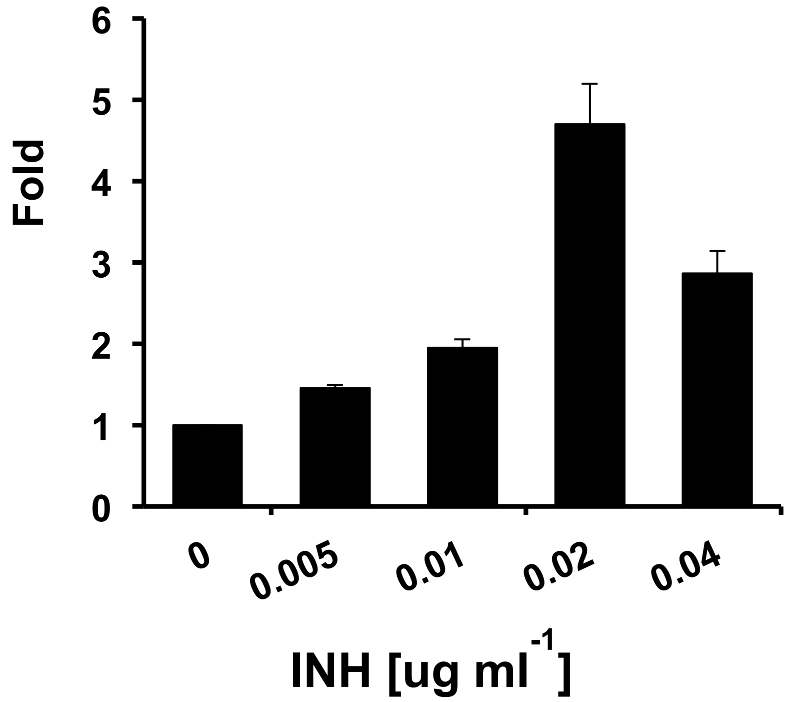


**Supplementary Figure 7. INH dependent *iniA* induction.** Induction of *iniA* in *Mtb* cells treated with increasing concentrations of INH in Fig. 5A, determined by RT-PCR. Fold change was calculated relative to untreated cells.

### Supplementary Table 1

**Plasmids used in this study**

| Plasmids | Characteristic(s) | Source/Ref |
|---|---|---|
| pSM316 | Integrative vector (Strp Spec Kan) | (42) |
| pSM986 | *iniBAC* gene cluster and its native promoter (-211bp of *iniB*) cloned into pSM316 | This study |
| pMV361 | Integrative vector for gene expression driven by hsp60 promoter | This study |
| pSM988 | *iniA* gene cloned into pMV361 | This study |
| pSM989 | *iniC* gene cloned into pMV361 | This study |

***M. tuberculosis* strains used in this study**

| Strains | Characteristic(s) | Source |
|---|---|---|
| H37Rv | Wild-type strain | ATCC |
| ST403 | *virR*::*tn*; Km$^{R}$ | (9) |
| ST421 | *iniA*::*hyg*; Hyg$^{R}$ | (23) |
| ST422 | ST421 harboring pSM986 | This study |
| ST424 | ST421 harboring pSM988 | This study |
| ST425 | ST421 harboring pSM989 | This study |

**Primer used in this study**

| Primer name | Sequence (5' to 3') | Application |
|---|---|---|
| *iniBAC*$_{\mathrm{TB}}$ com F | AATCTAGATGGGAGGCTTGATCATCACGGCTACGAC | *iniBAC* |
| *iniBAC*$_{\mathrm{TB}}$ com R | AATATTTCGCGAAGGCCTCAGCGGCGAGCAGAGAAC | complementation |
| *iniA*$_{\mathrm{TB}}$ com F | TCGCAATGATGGTCCCCGCCGGTTTG | |
| *iniA*$_{\mathrm{TB}}$ com R | GTCTTGGCTCACGCTCGTCCCAAGCTC | |
| pMV361_*iniA*$_{\mathrm{TB}}$ F | GAGCGTGAGCCAAGACAATTGCGGATCC | *iniA* |
| pMV361_*iniA*$_{\mathrm{TB}}$ R | GTCTTGGCTCACGCTCGTCCCAAGCTC | complementation |
| *iniC*$_{\mathrm{TB}}$ com F | TCGCAATGGTGAGCACCAGCGACCGGGTCCGC | |
| *iniC*$_{\mathrm{TB}}$ com R | GTCTTGGCTCAGCGGCGAGCAGAGAACTCCGC | *iniC* |
| pMV361_*iniC*$_{\mathrm{TB}}$ F | GCCGCTGAGCCAAGACAATTGCGGATCC | complementation |
| pMV361_*iniC*$_{\mathrm{TB}}$ R | GTGCTCACCATTGCGAAGTGATTCCTCC | |
| *iniA*$_{\mathrm{TB}}$ F | AAGATGATCCAGCGTCTGCT | RT-PCR |
| *iniA*$_{\mathrm{TB}}$ R | TTGACCTGGCTCAGGATACC | RT-PCR |

| | | |
|---|---|---|
| *P1-rrnA* $_{TB}$ F | CCTATGGATATCTATGGATGACCGA | RT-PCR |
| *P1-rrnA* $_{TB}$ R | GGCGACCCTGCCAGTCTAA | RT-PCR |
| Rv2628 F | AGGAGGCGATGATGAATCTAGC | RT-PCR |
| Rv2628 R | CAACTCCGACAACCAACCG | RT-PCR |
| *cmtR* (Rv1994c) F | AACCATCTGTCGTGTTTGCG | RT-PCR |
| *cmtR* (Rv1994c) R | ACACAGGGTTGGTCGGTATC | RT-PCR |
| *csoR* (Rv0967) F | GCAAGGAATTGACCGCAAAG | RT-PCR |
| *csoR* (Rv0967) R | AAGTGGTTGTGCAGCATCAC | RT-PCR |
| *ctpF* (Rv1997) F | TGAGATTCATCGGCTCGTTG | RT-PCR |
| *ctpF* (Rv1997) R | AACGTTTCGACGGCATCTTG | RT-PCR |
| *kmtR* (Rv0827c) F | ATGTACGCAGATAGTGGACCTG | RT-PCR |
| *kmtR* (Rv0827c) R | ATTCGTAGCTTTGCCAGGTG | RT-PCR |
| *lat* (Rv3290c) F | ACGACATCGCGTGTTTTGTG | RT-PCR |
| *lat* (Rv3290c) R | ACATCCAACTGCTGGTATGC | RT-PCR |
| *cysD* (Rv1285) F | AATCCGATACAGACCGTGACG | RT-PCR |
| *cysD* (Rv1285) R | TTGTAGAGGTTCCACAGTTCCG | RT-PCR |
| *cydB* (Rv1622c) F | ATTGTGGTTCGGTGTCATCG | RT-PCR |
| *cydB* (Rv1622c) R | GTGATCAGCCAGACTTCGTTG | RT-PCR |
| *acg* (Rv2032) F | GCATACCGCACAGTTCATTGG | RT-PCR |
| *acg* (Rv2032) R | TCAAGCAATACGGCGGAAAG | RT-PCR |
| *ppsC* (Rv2933) F | GCCAAACGGGAAATGCTTTC | RT-PCR |
| *ppsC* (Rv2933) R | TCTTGCCCAGTTCGATGAAC | RT-PCR |
| *iniB* (Rv0341) F | CTGGGTTTTACTGCCGTGATTG | RT-PCR |
| *iniB* (Rv0341) R | AACTCCGATCTGGCCATTGC | RT-PCR |

| | | |
|---|---|---|
| Rv1460 F | AAAATCCCGGCGGTCTCTAC | RT-PCR |
| Rv1460 R | ATCAGCGCGTCCAGATGAC | RT-PCR |
| Rv0324 F | AATATCCCGATAGCCGAACTGG | RT-PCR |
| Rv0324 R | CATTCGAGCATTCCGTCGTC | RT-PCR |
| *narK2* (Rv1737c) F | TCGTGATGCACCCTACTTTCG | RT-PCR |
| *narK2* (Rv1737c) R | AACCCGTAGATCGTGGTGATG | RT-PCR |
| hrp1 (Rv2626c) F | TCTACTACGTCGATGCGAACG | RT-PCR |
| hrp1 (Rv2626c) R | TTGACGAACTGCACAATGGC | RT-PCR |
| *dacB2* (Rv2911) F | AACAGCTGATCGTCAACCAG | RT-PCR |
| *dacB2* (Rv2911) R | TCTTTGACCAGCCCGTACATC | RT-PCR |
| *tgs1* (Rv3130c) F | AACAGGTGTGCCGGAAATTC | RT-PCR |
| *tgs1* (Rv3130c) R | TCTTGCTCAAAGCGCTGTTG | RT-PCR |
| *mctB* (Rv1698) F | ATCTCGTTGCGTCAACATGC | RT-PCR |
| *mctB* (Rv1698) R | AGCTTTTCGCGAAGTGCATC | RT-PCR |
